# sweetD: An R package using Hoeffding’s D statistic to visualise the dependence between M and A for large numbers of gene expression samples

**DOI:** 10.1101/2021.02.24.432640

**Authors:** Amber Barton

## Abstract

**Summary:** MA plots are frequently used to examine the relationship between gene abundance and differences in gene expression between two samples. In good quality samples without batch effects or outliers, there is generally no or little relationship between intensity and log fold difference. As the number of MA plots increases quadratically with the number of samples, for large datasets the number of potential MA plots becomes prohibitively high for manual examination. Here we present an R package calculating and visualising the dependence between abundance and log fold difference for large numbers of samples using Hoeffding’s D statistic.

**Availability and implementation:** sweetD is currently available as an R package on Github https://github.com/amberjoybarton/sweetD.

## 1 Introduction

MA plots are a common method of visualising the relationship between log ratio (M) and mean abundance (A) in gene expression data (McDermaid *et al*., 2019). The log ratio may be between one sample and average expression, between two samples, or between two groups. In the quality control steps of a differential expression pipeline, a relationship between M and A can imply batch effects, outlying samples or unsuccessful normalisation. As the number of samples in a gene expression dataset increases, the number of MA plots for sample-sample comparisons increases quadratically, and the number of MA plots for sample-median comparisons increases linearly. Therefore for large datasets it quickly becomes unfeasible to manually examine all MA plots. Furthermore, as the relationship between M and A can be non-monotonic, linear modelling or a correlation analysis is not suitable. However the non-parametric measure Hoeffding’s D statistic is able to assess dependence between two variables even in non-monotonic relationships (Hoeffding, 1948).

While Hoeffding’s D statistic has previously been used as a summary statistic for MA plots in the R package arrayQualityMetrics (Kauffmann *et al*., 2009), this package is limited in that expression data must be in a Bioconductor microarray dataset container, samples are compared with the median only, and only a small subset of samples are visualised. Here we present the R package sweetD, which calculates Hoeffding’s D statistic for all samples relative to the median or each other, which can take any log transformed gene expression matrix as an input, and which can simultaneously visualise changes in the distribution of Hoeffding’s D statistic over the course of a pre-processing pipeline.

## 2 Materials and Methods

sweetD provides five functions for calculation and visualisation of Hoeffding’s D statistic. sweetDmedian() and sweetDall() both take a series of log-transformed matrices as an input, with columns corresponding to samples and rows corresponding to genes or other features. sweetDmedian() calculates the median expression for each feature using the R stats package, then loops through samples calculating M and A compared with the median. sweetDall() on the other hand calculates M and A for each sample compared with every other sample. sweetDall() can be computationally intensive for large datasets and therefore users may benefit by limiting the analysis to a random subset of several thousand genes. The hoeffd() function in the Hmisc package is then used to calculate Hoeffding’s D statistic. For sweetDmedian() the output is a dataframe displaying the D statistic for each sample and input matrix, allowing users to compare D statistics for the same sample at different stages of a processing pipeline. For sweetDall() the output is a dataframe displaying the D statistic for each combination of samples and input matrix.

The outputs of sweetDmedian() and sweetDall() can then be input into the functions sweetDplot() and sweetDheatmap() respectively for visualisation. Both functions use ggplot2: sweetDplot() to display the distribution of D statistics for each input matrix, and sweetDheatmap() to display a heatmap of D statistics for every combination of samples in each input matrix.

The MAplot() function takes one log-transformed matrix and the names of two samples as an input. If only one sample is specified it is compared with the median. As above, M, A and Hoeffding’s D statistic are calculated. ggplot2 is used to directly display the relationship between M and A with Hoeffding’s D statistic displayed above the plot.

## 3 Case study

Publicly available dataset GSE30119, containing the transcriptional profiles of healthy controls (n = 44) and patients with acute *Staphylococcus aureus* infections (n = 99), was downloaded using the GEOquery getGEO() function in R (Banchereau *et al*., 2012; Sean and Meltzer, 2007). Data was log-transformed and either standardised using the preProcess() function in the caret package, or quantile normalised using the normalize.quantiles() function in the preprocessCore package (Kuhn, 2008).

Application of the sweetDmedian() and sweetDplot() functions from the sweetD package showed that 11/143 samples in the raw dataset had a Hoeffding’s D statistic > 0.1 relative to the median, compared with 9/143 following standardisation and 0/143 following quantile normalisation (Figure 1). The sample with the highest D statistic compared with the median in the raw and standardised data was GSM745758, a female patient with *Staphylococcus aureus* bacteraemia. Healthy controls were not over- or under- represented amongst samples with D > 0.1 in the raw dataset (27% in samples with D > 0.1 versus 31% across the entire dataset) suggesting that high D statistics were due to technical rather than biological variation. Furthermore, quantile normalisation appears to have been more successful than standardisation in accounting for this technical variation.

**Figure 1:**
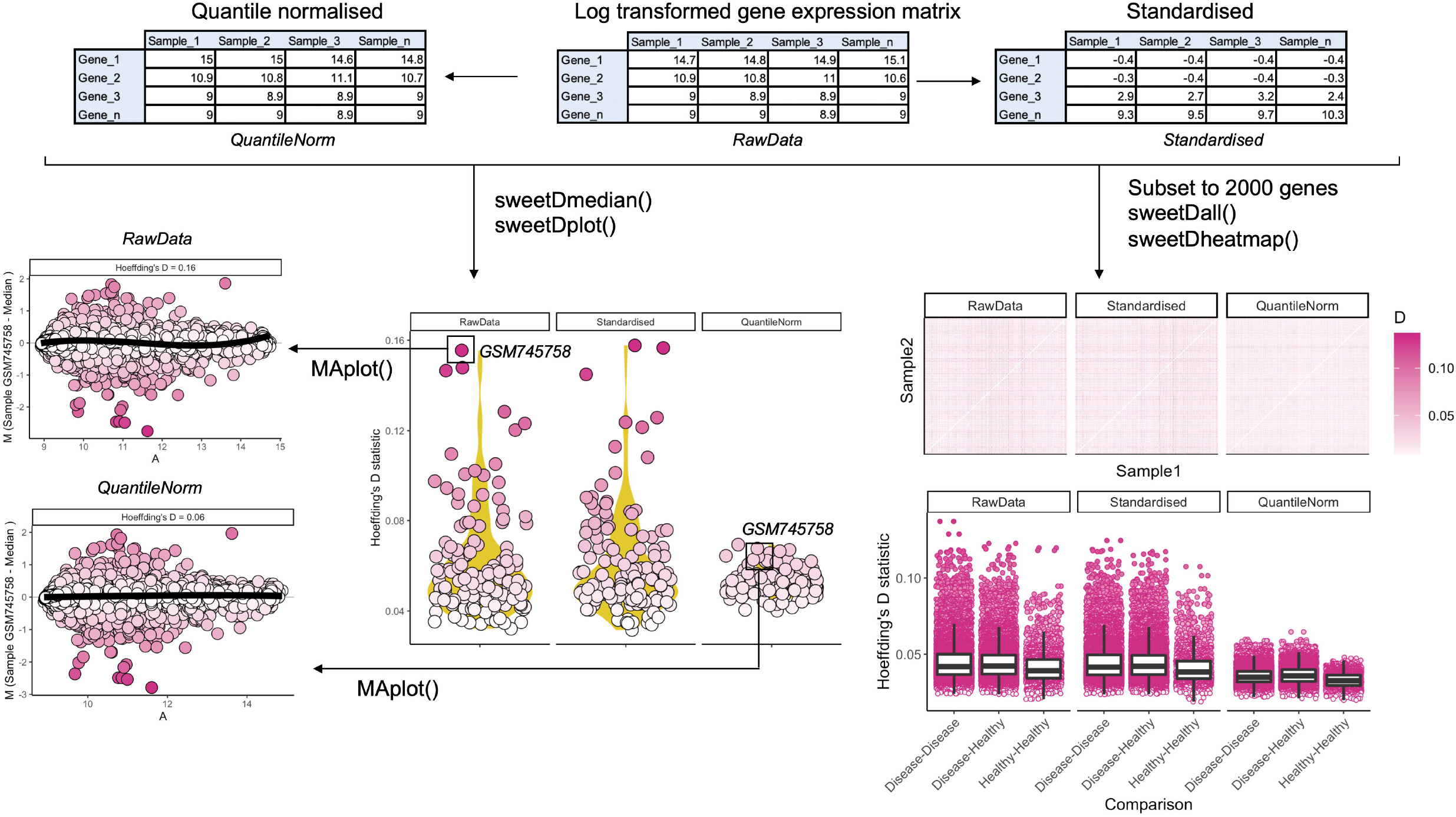
Publicly available dataset GSE30119 was log transformed then standardised or quantile normalised. Functions sweetDmedian() and sweetDplot() identified sample GSM745758 as having the highest Hoeffding’s D statistic relative to the median in the raw data. However this was brought down to D = 0.06 by quantile normalisation. Functions sweetDall() and sweetDheatmap() found that quantile normalisation brought down the number of sample-sample comparisons with a Hoeffding’s D statistic > 0.1. Comparisons between disease samples or between disease and healthy samples gave on average higher Hoeffding’s D statistics than comparisons between healthy samples.

The gene expression datasets were then subset to a random sample of 2000 genes and the functions sweetDall() and sweetDheatmap() applied to examine sample-sample Hoeffding’s D statistics. In this case, 96/20,449 sample-sample comparisons gave D > 0.1 in the raw dataset, 78/20,449 in the standardised dataset and 0/20,449 in the quantile normalised dataset. In all datasets Disease-Disease and Disease-Healthy comparisons overall yielded greater D statistics than Healthy-Healthy sample comparisons, suggesting greater biological variability amongst samples from those with *S. aureus* infection.

## 4 Conclusion

This article introduces the R package sweetD, allowing users to easily assess the dependence between log fold expression and abundance in gene expression data, and therefore identify outliers, batch effects and appropriate normalisation methods.

## Data Availability Statement

No new data were generated in support of this research. The dataset used in the case study is available in NCBI Geo, at https://www.ncbi.nlm.nih.gov/geo/query/acc.cgi?acc=gse30119.

## Notes

https://github.com/amberjoybarton/sweetD

### Competing Interest Statement

The authors have declared no competing interest.

